# The AP-1 complex regulates AXL expression and determines sensitivity to PI3Kα inhibition in esophagus and head and neck squamous cell carcinoma

**DOI:** 10.1101/415752

**Authors:** Mai Bdarny, Manu Prasad, Noa Balaban, Joshua Ben-Zion, Anat Bahat Dinur, Reidar Grénman, Limor Cohen, Moshe Elkabets

## Abstract

AXL overexpression is a common resistance mechanism to anti-cancer therapies, including the resistance to BYL719 (Alpelisib) – the p110α isoform specific inhibitor of phosphoinositide 3-kinase (PI3K) – in esophagus and head and neck squamous cell carcinoma (ESCC, HNSCC respectively). However, the mechanisms underlying AXL overexpression in resistance to BYL719 remain elusive. Here we demonstrated that the AP-1 transcription factors, c-JUN and c-FOS, regulate AXL overexpression in HNSCC and ESCC. The expression of AXL was correlated with that of c-JUN both in HNSCC patients and in HNSCC and ESCC cell lines. Silencing of c-JUN and c-FOS expression in tumor cells downregulated AXL expression and enhanced the sensitivity of human papilloma virus positive (HPV^Pos^) and negative (HPV^Neg^) tumor cells to BYL719 in vitro. Blocking of the c-JUN N-terminal kinase (JNK) using SP600125 in combination with BYL719 showed a synergistic anti-proliferative effect in vitro, which was accompanied by AXL downregulation and potent inhibition of the mTOR pathway. In vivo, the BYL719-SP600125 drug combination led to the arrest of tumor growth in cell line-derived and patient-derived xenograft models, and in syngeneic head and neck murine cancer models. Collectively, our data suggests that JNK inhibition in combination with anti-PI3K therapy is a new therapeutic strategy that should be tested in HPV^Pos^ and HPV^Neg^ HNSCC and ESCC patients.

## Introduction

The unsatisfactory outcomes of current therapies for patients with squamous cell carcinoma of the head and neck (HNSCC) or of the esophagus (ESCC) are underscored by the high mortality and morbidity rates of these cancers (1, 2). These outcomes, taken together with the high incidence rates of HNSCC and ESCC in many countries (3, 4) indicate an urgent need for new therapeutic approaches.

The major causes for the development of HNSCC and ESCC are alcohol consumption, smoking, and infection with human papilloma virus (HPV) [reviewed in (5)]. Despite significant differences in the molecular landscapes of HPV-positive (HPV^Pos^) and HPV-negative (HPV^Neg^) HNSCC and ESCC (6–8), both are frequently associated with alterations in the *PIK3CA* gene (6–8), which encodes for the p110α subunit of the phosphatidylinositol-4,5-bisphosphate 3-kinase (PI3K). Activating point mutations or amplifications of the *PIK3CA* gene result in the hyper-activation of the PI3K/AKT/mammalian target of rapamycin (mTOR) signaling pathway [reviewed in (9)]. This pathway plays a key role in regulating cell proliferation and survival, enhancing tumor progression in PIK3CA-mutated HNSCC and ESCC. It is thus self-evident that new approaches for the treatment of HNSCC and ESCC should focus on blockers of the components of the PI3K/AKT/mTOR pathway, and indeed such blockers are now under clinical development [reviewed in (10–12)]. Among these blockers is the compound designated BYL719 (Alpelisib), which is an isoform-specific p110α inhibitor. In the first-in-human study of this compound, Juric et al. reported that 14 out of 17 patients with PIK3CA-mutated HNSCC benefited from single agent administration of BYL719 (13), although all patients eventually developed resistance to BYL719 (13). We recently showed that the emergence of resistance to BYL719 in HNSCC and ESCC involves the overexpression of AXL, which is a receptor tyrosine kinase (RTK) (14). AXL dimerizes with epithelial growth factor receptor (EGFR) to activate the phospholipase Cγ (PLCγ)-protein kinase C (PKC) signaling pathway, leading to the activation of mTOR in an AKT-independent manner (14). We further showed that AXL overexpression is associated with resistance to BYL719 in patients with HNSCC and that inhibition of AXL using R428 could reverse the resistance to BYL719 (14). Other studies have shown that AXL overexpression plays a key role in the resistance to many other anti-cancer therapies (15–19). These lines of evidence signify that treatment efficacies could be improved by blocking AXL activity, and, indeed, small-molecule and antibody blockers of AXL are currently under clinical trials (ClinicaTrials.gov). However, to the best of our knowledge, targeting the expression of AXL as an alternative therapeutic strategy – as will be described here – has not been explored to date.

AXL gene transcription has been shown to be regulated by several transcription factors (TFs), such as SP1/3 (20) and MZF1 (21) in colon and cervix cancers, and the AP-1 complex in melanoma, leukemia and bladder cancer (22–24). However, the TFs that regulate AXL overexpression in ESCC and HNSCC in resistance to PI3K therapy remain uncharacterized. Here, we sought to elucidate the transcriptional machinery that regulates AXL expression and to explore whether a treatment protocol targeting AXL transcription in combination with BYL719 could serve as a therapeutic opportunity in HNSCC and ESCC.

## Results

### AXL expression determines sensitivity to BYL719 in HPV^Pos^ and HPV^Neg^ cancer cell lines

We have recently shown that AXL overexpression drives the resistance to BYL719 in HNSCC and ESCC cell lines and in HNSCC patients (14). In the current study, we first examined whether the basal expression of AXL determines the primary sensitivity to BYL719 in HPV^Pos^ and HPV^Neg^ cell lines. As HPV^Neg^ cell lines, we used SNU1076 (an HNSCC cell line) and our previously established isogenic tumor cell line model, BYL719-sensitive KYSE180 (KYSE180^Sen^), and its counterpart BYL719-resistant model (KYSE180^Res^) that showed AXL overexpression (ESCC cell lines) (14). As the HPV^Pos^ cell line, we used UM-SCC-47. To this end, we knocked down the expression of AXL in these HPV^Pos^ and HPV^Neg^ HNSCC and ESCC cell lines, and measured the half maximal inhibitory concentration (IC:50) of BYL719 in vitro. Knockdown of AXL significantly reduced BYL719 IC:50 values in all tumor cells (Figure 1A). Western-blot (WB) analysis of the KYSE180^Sen^ tumor cells following BYL719 treatment showed that AXL expression impacted the ability of BYL719 to block the mTOR pathway. Specifically, intensified inhibition of the phosphorylation of ribosomal protein S6 (pS6) was detected in tumor cells in which AXL expression had been knocked down (Figure 1B). These results are in keeping with our previous findings that AXL bypasses AKT to activate the mTOR pathway (14). Next, we confirmed the effect of AXL knockdown on the efficacy of BYL719 in vivo in a cell line-derived xenograft model (CDX) of KYSE180^Sen^ tumor cells. In agreement with the above in vitro studies, BYL719-induced inhibition of growth was more efficient in the AXL knocked-down tumor cells (Figure 1C). This growth inhibition was associated with a reduction in mTOR pathway activation [as evaluated by pS6 immunohistochemical (IHC) staining] and, consequently, with a reduction in tumor cell proliferation [as determined by Ki67 IHC staining] (Figure 1D and E). Similar results were obtained for the HPV^Pos^ UM-SCC-47 cells (Supplementary Figure 1A-D).

**Figure 1:**
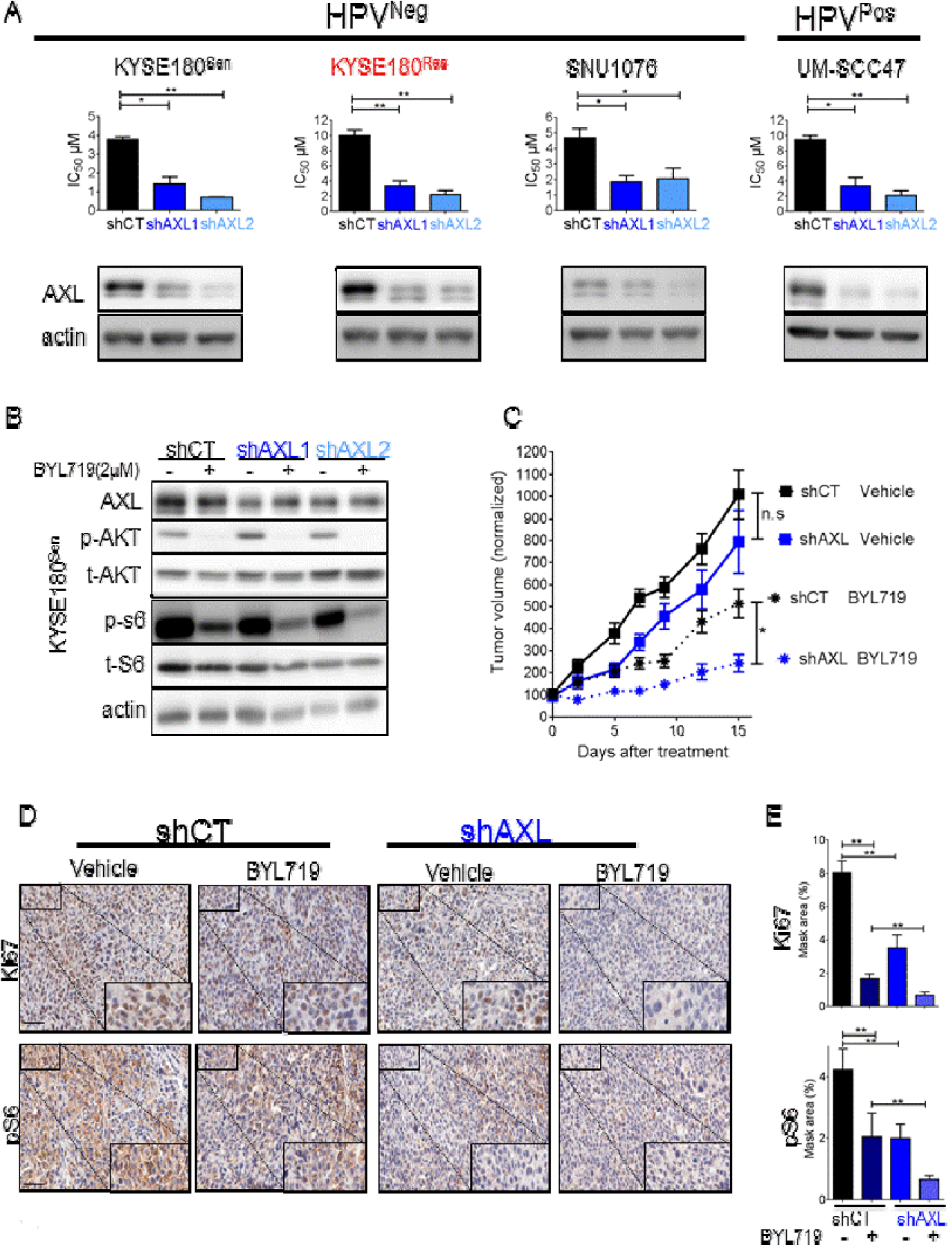
AXL knockdown sensitizes HNSCC and ESCC cells to BYL719 in vitro and in vivo. **A.** Upper-panel-AXL (shAXL1 and shAXL2) compared with control cells (shCT). Lower panel - WB analysis demonstrating AXL levels. **B.** WB analysis showing AKT/mTOR pathway activation in KYSE180^Sen^ cells after AXL knockdown and BYL719 treatment (2 μM, 24 hours). **C.** Tumor growth of shCT or shAXL1 KYSE180^Sen^ CDXs in mice treated daily with BYL719 25mg/kg. **D.** IHC analysis of the tumors showing the expression levels of the proliferation marker Ki67 and phosphorylated ribosomal S6 levels (pS6). The size of the scale bar represents 50μm. A higher magnification is shown in the insert. **E.** Analysis of expression levels of IHC staining presented in D, using the 3DHISTECH software HistoQuant™. *P = 0.05; **P= 0.01.

### Resistance to BYL719 is associated with upregulation of the c-JUN transcription factor

Since we had shown that the expression level of AXL determined the response of HNSCC and ESCC cells to BYL719, we aimed to identify the molecular machinery that regulates AXL overexpression in our previously reported isogenic BYL719-sensitive and -resistant models, namely, KYSE180^Sen^ vs. KYSE180^Res^ and CAL33^Sen^ vs. CAL33^Res^ tumor cell lines (14). A gene set enrichment analysis of the RNA-seq data obtained for these cells identified over 100 conserved motifs among 50 TFs that were significantly activated (Supplementary Table 1). TCF3, SP-1, AP-1 and MYC were among the most significant TF signatures that were upregulated in both resistant cell lines. Of the 50 TFs, three were predicted to have binding sites on the AXL promotor (QIAGEN transcription factor analysis), namely, AP-1, MYC, and MYC-associated zinc finger protein (MAZ) (Figure 2A). Further analysis of the RNA-seq data indicated that the expression levels of MAZ and MYC were similar in BYL719-sensitive and BYL719-resistant cells, whereas the genes of the AP-1 complex, namely, FOSL1, FOS and c-JUN, were significantly upregulated in both resistant cell models (Supplementary Figure 2A). WB and qPCR analyses confirmed the upregulation of c-JUN in BYL719-resistant vs. sensitive cells, with c-JUN upregulation being associated with AXL overexpression (Figure 2B and C). The SP1 TF [previously reported to regulate AXL expression (20, 25)] was upregulated in KYSE180^Res^ cells but not in CAL33^Res^ cells. In addition, we found an inverse correlation between AXL and the microphthalmia-associated transcription factor (MITF) (Figure 2B), in agreement with previous reports (23, 26, 27). Increased expression of AXL and c-JUN was further confirmed in SNU1076 cells that had acquired resistance to BYL719 (Supplementary Figure 2B).

**Figure 2:**
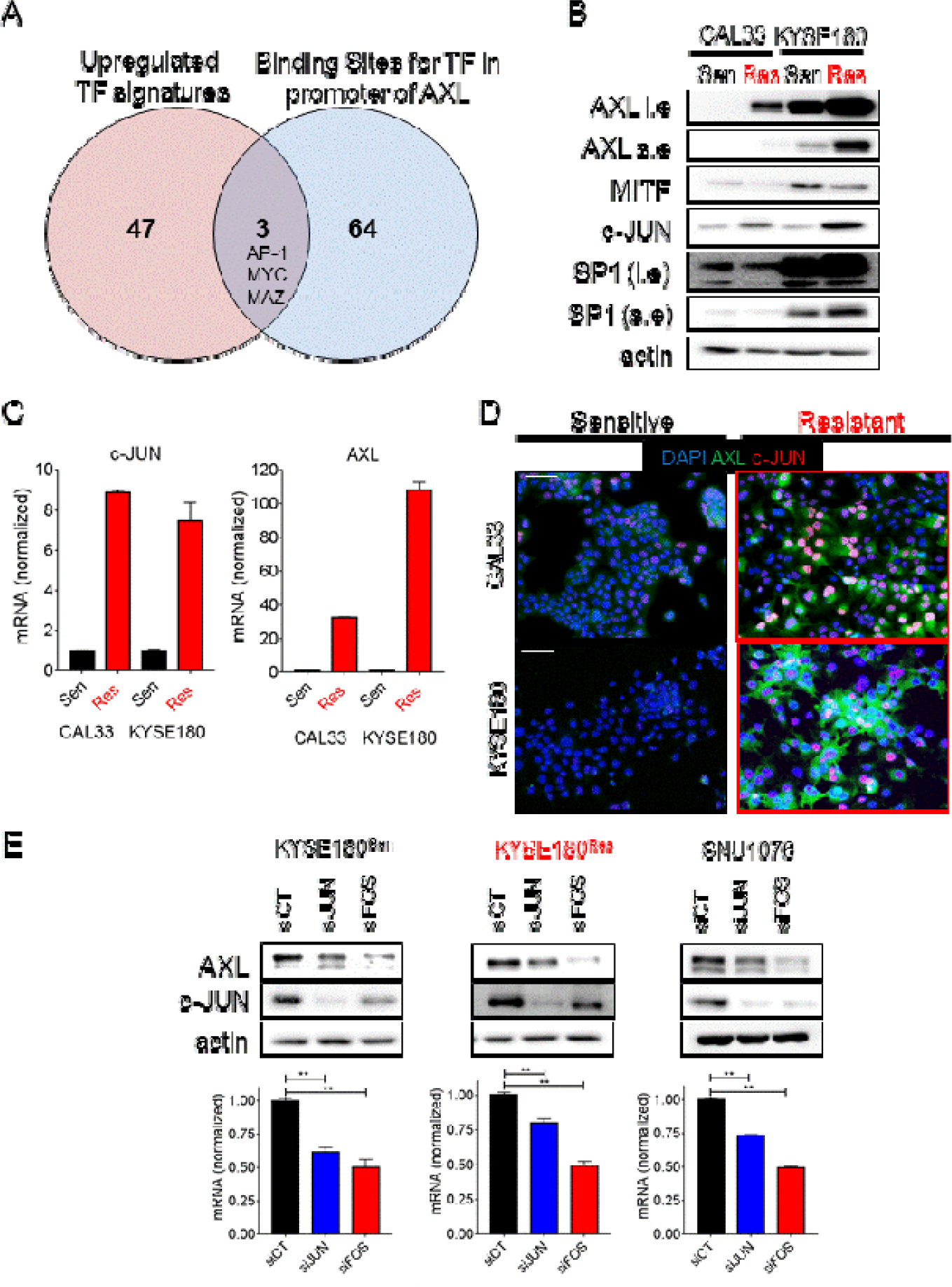
The AP-1 transcriptional complex regulates AXL expression in HNSCC and ESCC. **A.** RNA sequencing data obtained from isogenic BYL719 sensitive vs. acquired-resistance CAL33 and KYSE180 cells (HNSCC and ESCC, respectively). Presented in the Venn diagram are TF signatures upregulated in BYL719 resistant cells and their overlap with binding sites for TF in the promoter of AXL. **B.** WB analysis of CAL33 and KYSE180 BYL719-sensitive vs. resistant cells showing the expression of TFs (l.e - long exposure; s.e - short exposure) **C.** A qPCR analysis comparing the mRNA levels of AXL and c-JUN in CAL33 and KYSE180 sensitive vs. resistant cells. **D.** IF images of CAL33 and KYSE180 BYL719-sensitive vs. resistant cells showing c-JUN (CY-3 labeled) in red, AXL (Alexa488 labeled) in green and DAPI in blue. The size of the scale bar represents 100μm. **E.** Upper panel - WB analysis showing AXL levels after transfection with siRNA for the silencing of c-JUN and c-FOS. Lower panel - qPCR analysis showing AXL mRNA levels in cells transfected with siRNAs for the silencing of c-JUN and c-FOS.

To obtain further insight into AXL and c-JUN expression at the single cell level, immunofluorescent (IF) staining and imaging flow cytometry were performed. IF staining of AXL and c-JUN showed a correlation between the upregulation of AXL and that of c-JUN in BYL719-resistant cells at the single cell level (cells with high AXL displayed high c-JUN levels) (Figure 2D). Imaging flow cytometry analysis demonstrated that 97% of the KYSE180^Sen^ cells displayed low expression levels of both c-JUN and AXL, and only a small subset of the tumor cells showed high expression levels of c-JUN that were accompanied by high AXL expression. This trend was reversed in KYSE180^Res^ cells, most which (96.2%) showed high expression levels of both c-JUN and AXL (Supplementary Figure 2C). Given the positive correlation between AXL and c-JUN, we posited that the AP-1 transcriptional complex, which includes c-JUN and its counterpart c-FOS, regulates AXL expression. Indeed, silencing of c-JUN or c-FOS expression led to downregulation of AXL in KYSE180^Sen^, KYSE180^Res^, and SNU1076 cells, as evaluated by WB and qPCR analysis (Figure 2E and Supplementary Figure 2D and E).

### AXL and c-JUN levels are correlated in clinical samples from HNSCC patients and in HNSCC and ESCC cell lines

To confirm the role played by c-JUN in AXL expression in HNSCC, we tested (using IHC staining) the correlation between AXL and c-JUN levels in patients with malignant HNSCC (n = 17) and in those with benign head and neck tumors (n = 15). IHC staining showed that AXL and c-JUN were positively correlated, both in tissues from the HNSCC patients (R = 0.450, p = 0.0051, Figure 3A) and in benign tissues (R = 0.6772, p <0.0001, Supplementary Figure 3B and Supplementary Table 1). Moreover, our finding of a significant increase in both AXL and c-JUN expression in HNSCC compared to the benign tissues (Supplementary Figure 3A) is in keeping with previous reports of elevated c-JUN expression in oral SCC (28) (Supplementary Figure 3C). Validation of the expression of AXL and c-JUN was performed in an independent group of 10 freshly isolated HNSCC tumor specimens obtained from Soroka Medical Center; the samples analyzed by WB and IHC showed a similar trend for the association of the expression levels of AXL and c-JUN (R = 0.5254, p = 0.011, Figure 3B). In addition, a positive correlation between AXL and c-JUN expression was detected in 5 HNSCC patient-derived xenografts (PDXs) (R = 0.969, p = 0.0063, Supplementary Figure 3D) and in a panel of 17 HNSCC and ESCC cell lines (R = 0.308, p = 0.0085, Figure 3C). The BYL719 IC:50 values for the tested HNSCC and ESCC cell lines were also correlated to AXL and c-JUN expression, with most cell lines having high IC:50 values also showing high expression levels of AXL and c-JUN (shown by the sizes of the round symbols in the plots representing each cell line in Figure 3C and in Supplementary Figure 3E). Collectively, the strong positive correlation found between AXL and c-JUN expression in HNSCC and ESCC clinical samples and in cell lines supports our premise that AXL expression is transcriptionally regulated by the AP-1 complex.

**Figure 3.**
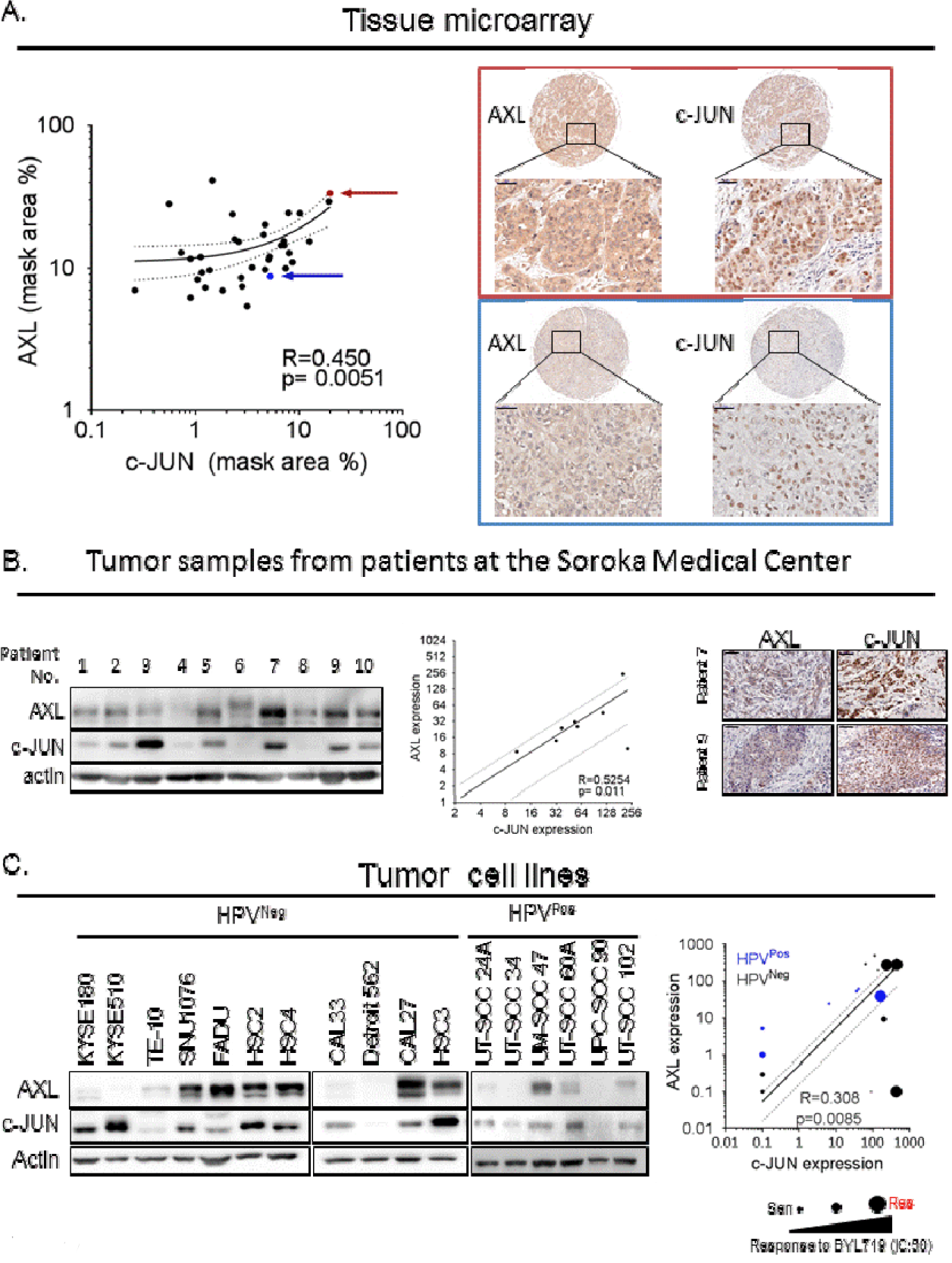
AXL and c-JUN levels are correlated in clinical samples of HNSCC tumors and in cell lines. **A.** IHC analysis of AXL and c-JUN expression levels in a tissue array of HNSCC tumors. The brown and blue arrows indicate two different tumor samples that correlate with the IHC images marked by the brown and blue rectangles on the left. AXL and c-JUN expression levels were calculated using the 3DHISTECH software HistoQuant™. The size of the scale bar represents 50μm. **B.** WB analysis of AXL and c-JUN expression levels in patient tumor samples from Soroka Medical Center. A densitometry analysis of expression levels of AXL and c-JUN is presented in the graph. Left - IHC images of tumor tissues demonstrating AXL and c-JUN expression. The size of the scale bar represents 50μm. **C.** WB analysis of HPV^Neg^ and HPV^Pos^ HNSCC and ESCC cell lines, demonstrating the expression levels of AXL and c-JUN. The graph on the right shows quantification of the WB analysis. The size of the symbol correlates to the BYL719 IC:50 value of the cell line in μM (i.e, the larger the symbol, the higher the IC:50), and HPV status represented by a black for HPV^Neg^ or blue forHPV^Pos^ symbol, IC:50 values for HPV^Neg^ cells were extracted from our previous report (14).

### Knockdown of members of the AP-1 complex and inhibition of the c-JUN N-terminal kinase enhance BYL719 efficacy in vitro

As AXL expression was shown to be regulated by the AP-1 transcription complex, we posited that knockdown of c-JUN and c-FOS would sensitize HNSCC and ESCC cells to BYL719. Indeed, we found that silencing of c-JUN and c-FOS expression in HPV^Neg^, KYSE180^Sen^, KYSE180^Res^ and SNU1076 cells significantly reduced the IC:50 values of BYL719. Similar results were obtained in the HPV^Pos^ cell lines UT-SCC-60A and UM-SCC-47 (Figure 4A and Supplementary Figure 4A).

**Figure 4:**
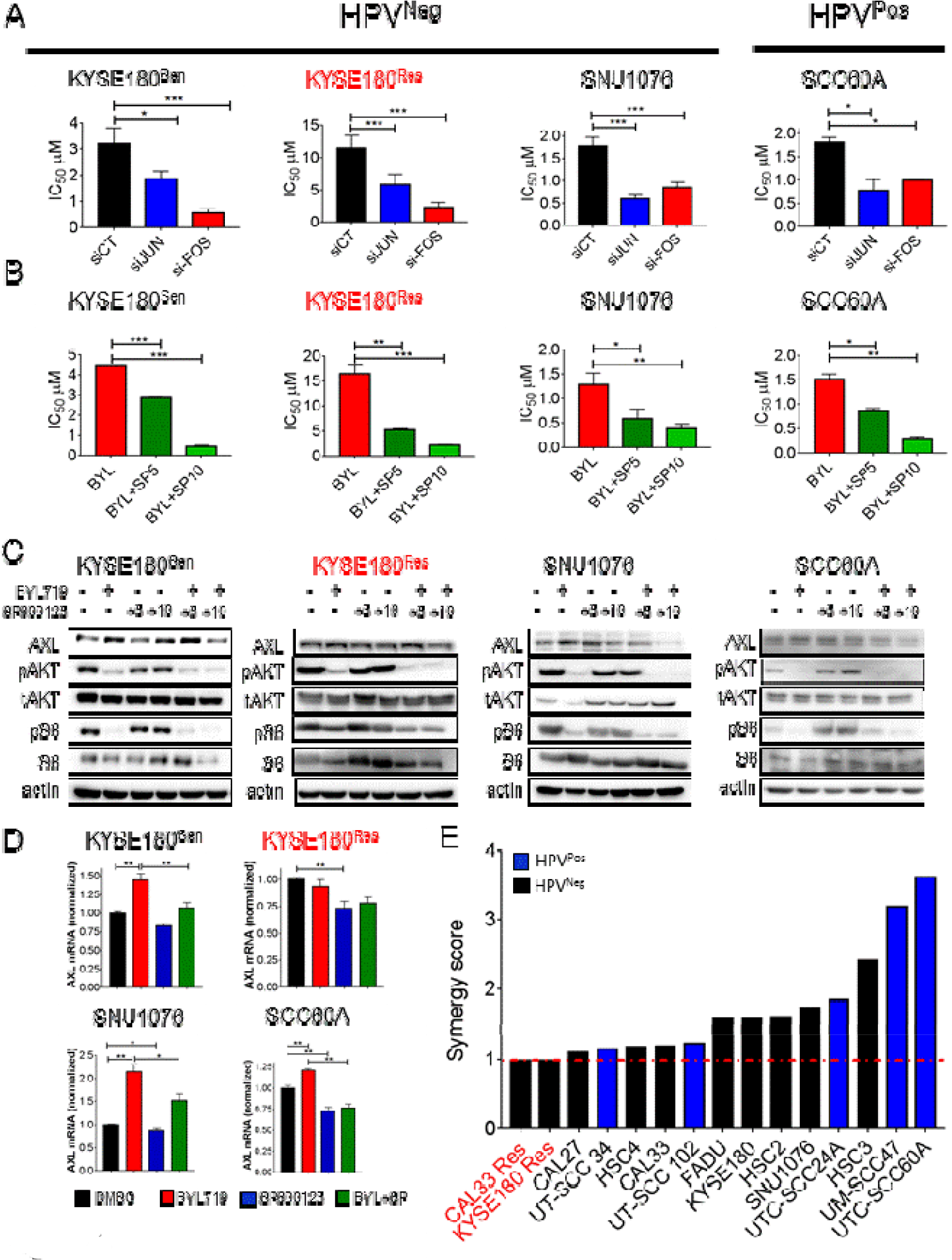
Silencing of c-JUN and c-FOS or blocking c-JUN N-terminal kinase (JNK) sensitizes HNSCC and ESCC cells to BYL719 in vitro. **A.** Analysis of BYL719 IC:50 values in HNSCC and ESCC cells following transfection with siRNAs to silence c-JUN and c-FOS expression. **B.** Analysis of BYL719 IC:50 values following JNK inhibition with SP600125 (5 and 10 μM) in HNSCC and ESCC cells. **C.** WB analysis showing AXL level, and AKT/mTOR pathway activation in HNSSC and ESCC cells treated with BYL719 (2 μM), SP600125 (5 and 10 μM) and the combination therapy for 24 hours. **D.** qPCR analysis of AXL mRNA levels in cells treated as in with BYL719 (2μM), SP600125 (10 μM) and combination for 24 hours. **E.** Synergy test for the interaction between BYL719 and SP600125. The synergy test was analyzed using Chalice software (Horizon), and a synergy score was extracted. *P = 0.05; **P= 0.01.

The direct role of c-JUN/c-FOS in AXL expression (Figure 2) and in the sensitivity to BYL719 (Figure 4A) suggests that blocking of c-JUN activity may potentiate BYL719 efficacy. As direct blockers of c-JUN are not yet available, we tested the above premise with an inhibitor of c-JUN N-terminal kinase (JNK), designated SP600125 (29). A combination of BYL719 and SP600125 significantly decreased the IC:50 values of BYL719 in three HPV^Neg^ and in two HPV^Pos^ HNSCC and ESCC cell lines in a dose dependent manner (Figure 4B and Supplementary Figure 4B). Notably, the IC:50 values for KYSE180^Res^ cells treated with the BYL719–SP600125 combination were similar to those for KYSE180^Sen^ cells, which suggests that the combined treatment delayed the acquisition of resistance. A cell proliferation assay monitoring growth rates of the cells showed that the combined BYL719–SP600125 treatment had a superior anti-proliferative effect compared to treatment with each of the compounds alone (Supplementary Figure 4C). WB analysis showed that while BYL719 upregulated AXL expression, its combination with SP600125 (10 μM) resulted in decreased AXL protein levels (Figure 4C). The level of pS6 was further inhibited by the combination of BYL719 and SP600125 in KYSE180^Res^ and SNU1076 cells (Figure 4C), when compared to the effect of BYL719. For KYSE180^Sen^, SNU1076 and UT-SCC-60A cells, qPCR analysis showed that AXL mRNA levels increased following BYL719 treatment but this gene upregulation was attenuated in response the combination with SP600125 (Figure 4D). In KYSE180^Res^ cells, in which AXL transcript basal levels were stably elevated (Figure 2C), a significant downregulation of AXL mRNA levels was detected following exposure to SP600125.

To explore whether the anti-proliferative effect of BYL719 and SP600125 is additive or synergistic, a synergy test was performed in five PIK3CA^mut^ and three PIK3CA^wt^ (all HPV^Neg^) HNSCC and ESCC cell lines, and five HPV^Pos^ HNSCC cell lines. Representative images of the analysis in SNU1076 cells are presented in Supplementary Figure 4D. Calculation of the synergy score using Chalice™ software indicated a synergistic effect (>1) in 13 of the 15 cell lines tested, whereas an additive effect was observed in the BYL719-acquired-resistant cell lines KYSE180^Res^ and CAL33^Res^ cell lines (Figure 4E).

### Superior efficacy of the combined treatment with BYL719 and SP600125 in vivo

To explore the efficacy of the BYL719–SP600125 combination in vivo, CDX models of KYSE180^Sen^ (HPV^Neg^) and UM-SCC-47 (HPV^Pos^) cells were generated. Daily treatments of tumor-bearing mice with BYL719 monotherapy (25 mg/kg) delayed tumor growth compared to treatment with the vehicle. Importantly, marked arrest of tumor growth (in HPV^Neg^ KYSE180^Sen^ CDXs) and significant tumor shrinkage (in HPV^Pos^-UM-SCC-47 CDXs) were observed in mice treated with the BYL719– SP600125 combination (Figure 5A and B, Supplementary Figure 5A and B). We did not detect any toxicity in that the weight of the animals remained stable (Supplementary Figure 5A and B). IHC staining of the proliferation marker, Ki67, showed decreased tumor cell proliferation in tumors treated with either BYL719 or SP600125 monotherapy. However, a more marked inhibition of cell proliferation was detected in tumors treated with the combination therapy.

**Figure 5:**
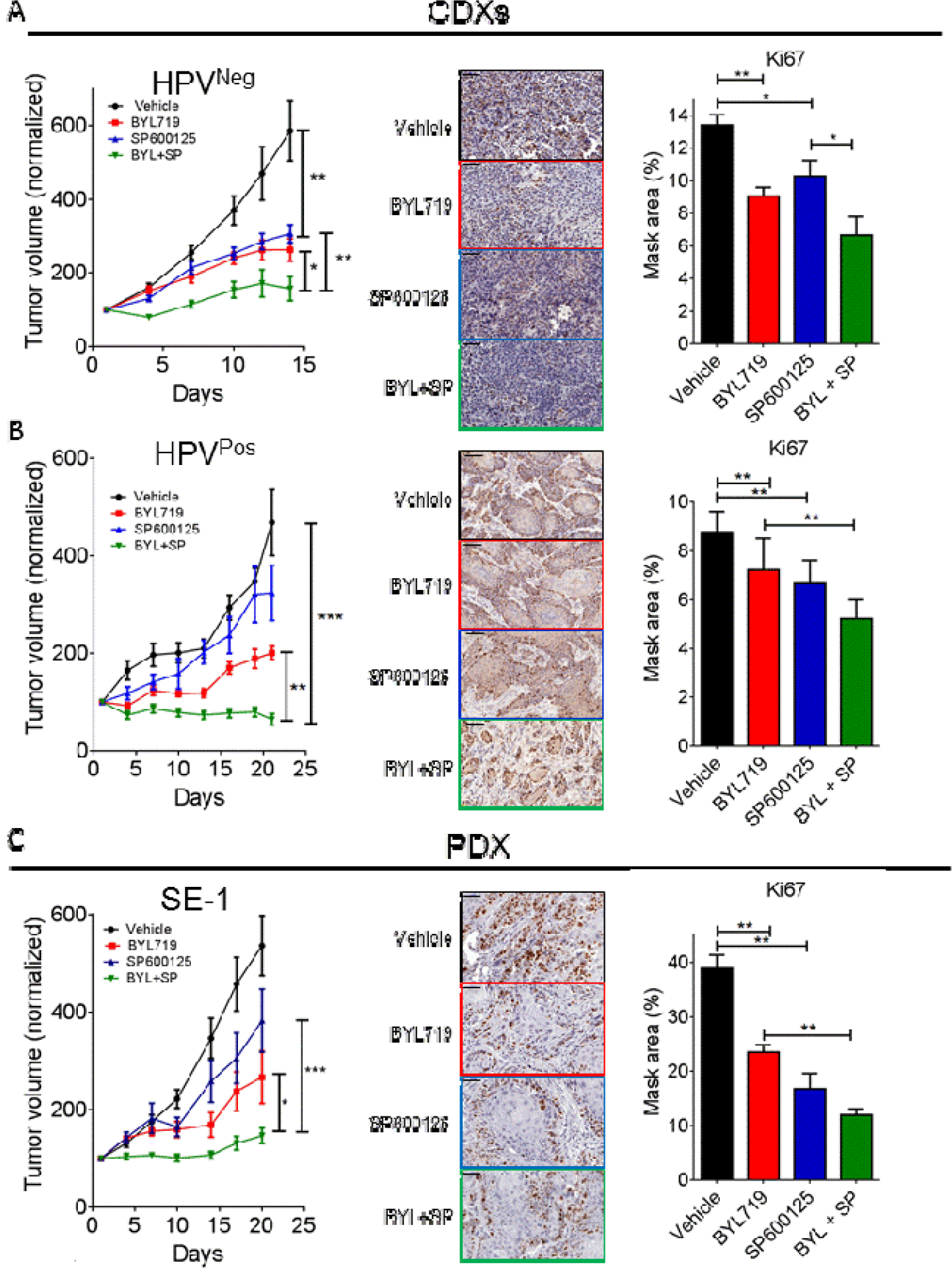
SP600125 enhances BYL719 efficacy in vivo in CDX and PDX models. **A-C**. Tumor growth of KYSE180^Sen^ **(A)**, UTSCC47 **(B)** and HNSCC PDX-SE1 **(C)** following daily treatment with vehicle, BYL719 (25 mg/kg), SP600125 (15 mg/kg) or the BYL719–SP600125 combination. IHC staining of the tumors for the proliferation marker Ki67 is shown on the left. The size of the scale bar represents 50μm. The 3DHISTECH software HistoQuant™ was used analysis of IHC staining. *P = 0.05; **P= 0.01; ***P = 0.001.

To further validate our observations in pre-clinical models, the efficacy of the drug combination was tested in two PDX models (SE1 and SE3). Treatment of the mice with the BYL719–SP600125 combination confirmed the superior anti-tumor effect of the combination, as indicated by smaller tumor volumes and lower tumor weights (Figure 5C and Supplementary Figure 5C). Ki67 staining showed a similar trend, further supporting the potency of drug combination.

### Enhanced anti-tumor efficacy of the BYL719–SP600125 combination prevents tumor progression and enhances survival in syngeneic head and neck cancers

To verify the efficacy of the combination of BLY719 and SP600125 in syngeneic HNSCC tumors, we generated HPV^Neg^ tumors in C57BL/6 mice. To this end, C57BL6/c mice were exposed to the carcinogen 4-nitroquinoline 1-oxide (4NQO) in their drinking water. Lip and tongue tumors developed following the exposure of the mice to 4NQO, as previously reported (30) (Supplementary Figure 6A). IHC staining demonstrated the activation of the AKT pathway, as well as the expression of c-JUN and AXL in the 4NQO-induced-tongue malignant lesion [Ki67 and KRT14 positive staining (Figure 6A)]. Following the generation of two cell lines derived from the lip and tongue tumors, we studied the efficacy of BYL719, SP600125 and the combination in vitro and in vivo. In vitro, the BYL719–SP600125 combination exhibited superior anti-proliferative activity, which was associated with a reduction of pS6 levels, as shown by WB analysis (Figure 6B and C). In vivo, the efficacy of the drug combination had a significant anti-tumor effect in the two syngeneic cancer models. Measurements of the lip tumors in mice treated with the BYL719–SP600125 combination indicated a stable disease, whereas tumor progression was observed in mice treated with either BYL719 or SP600125 (Figure 6E). In an orthotropic tongue cancer model, the BYL719–SP600125 combination significantly improved the survival of the mice. The median survival for vehicle- and SP600125-treated mice was 16 and 17 days, respectively, and for BYL719-treated mice, 23 days. Importantly, the median survival of mice receiving the combination therapy was more than doubled, being 52 days. MRI imaging supported the findings; the scans showed, in particular, the inhibition of tumor progression with the drug combination (Figure 6D).

**Figure 6:**
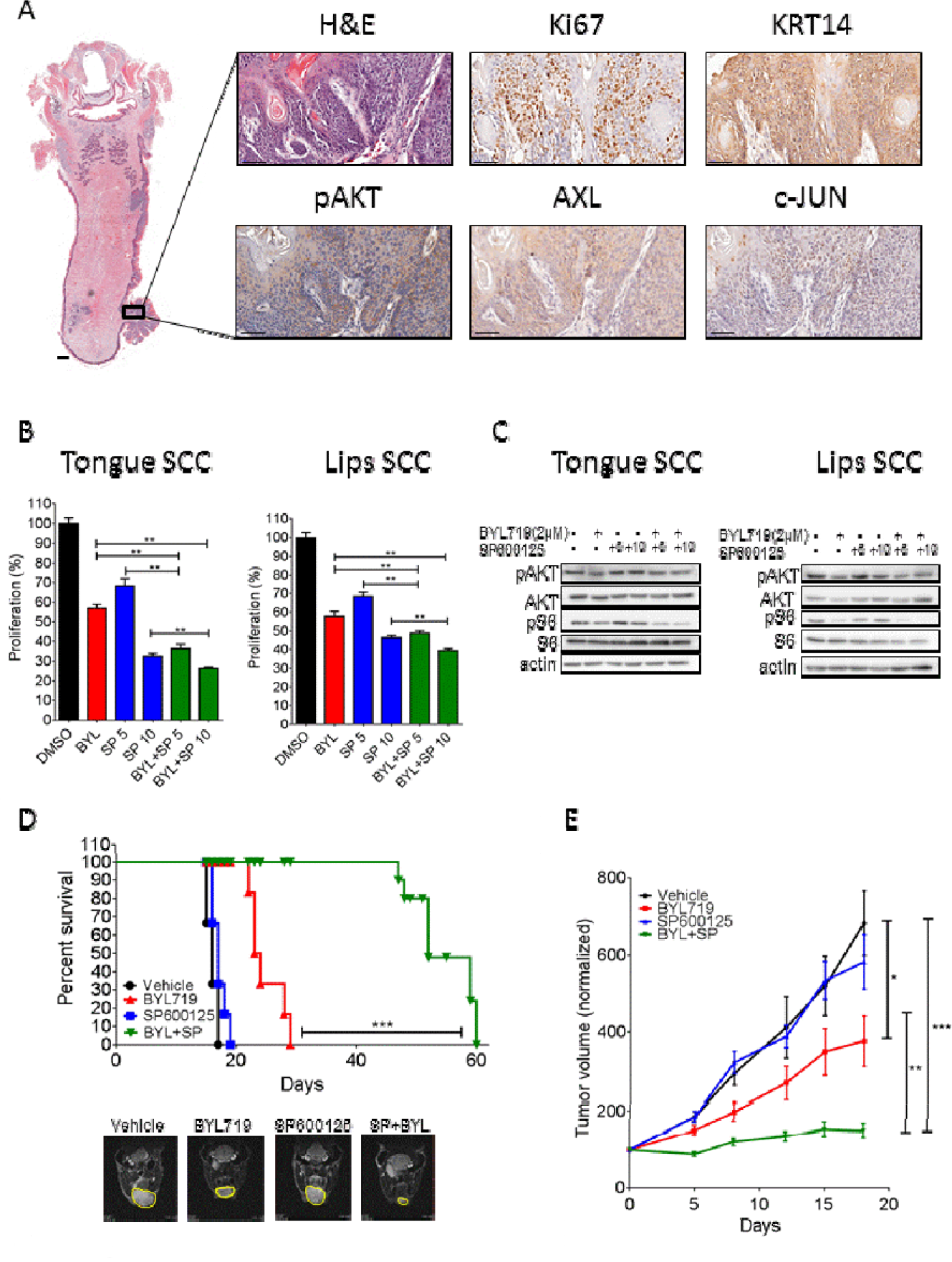
SP600125 increases BYL719 efficacy in vivo in syngeneic head and neck cancer models. **A.** IHC staining showing the expression of Ki67, pAKT, AXL, c-JUN and keratin-14 (KRT14) in the 4NQO-induced tongue tumor region. The size of the scale bars represents 1000μm in the small magnification image of whole tongue, and 50μm for the higher magnification images. **B**. Proliferation assay of 4NQO-induced tumor cell lines from the tongue and lips following 4 days of treatment with BYL719 (2 μM), SP600125 (5 and 10 μM) or the BYL719–SP600125 combination. **C.** WB analysis of tongue and lip 4NQO-induced tumor cell lines after 24 hours with the indicated treatments, showing activation of the AKT/mTOR pathway. **D.** Survival rates of immunocompetent C57BL/6 mice in an orthotopic tongue cancer model following daily treatments with BYL719 (25 mg/kg), SP600125 (15 mg/kg) or the BYL719–SP600125 combination. Also presented are T2 weighted coronal images (obtained from the MRI) of tongues (lower panel), showing the difference in tumor progression following 10 days of treatments. **E.** Tumor growth of immunocompetent C57BL/6 mice bearing a syngeneic lip cancer model, treated as indicated in D. **P= 0.01.

## Discussion

In this work, we demonstrated that the AXL expression level influences the sensitivity to BYL719 of HPV^Neg^ and HPV^Pos^ HNSCC and ESCC cells. We identified the AP-1 transcriptional complex as a regulator of AXL expression in these diseases and demonstrated that inhibition of JNK enhances the anti-tumor efficacy of BYL719 in vitro and in vivo (Scheme 1).

**Scheme 1:**
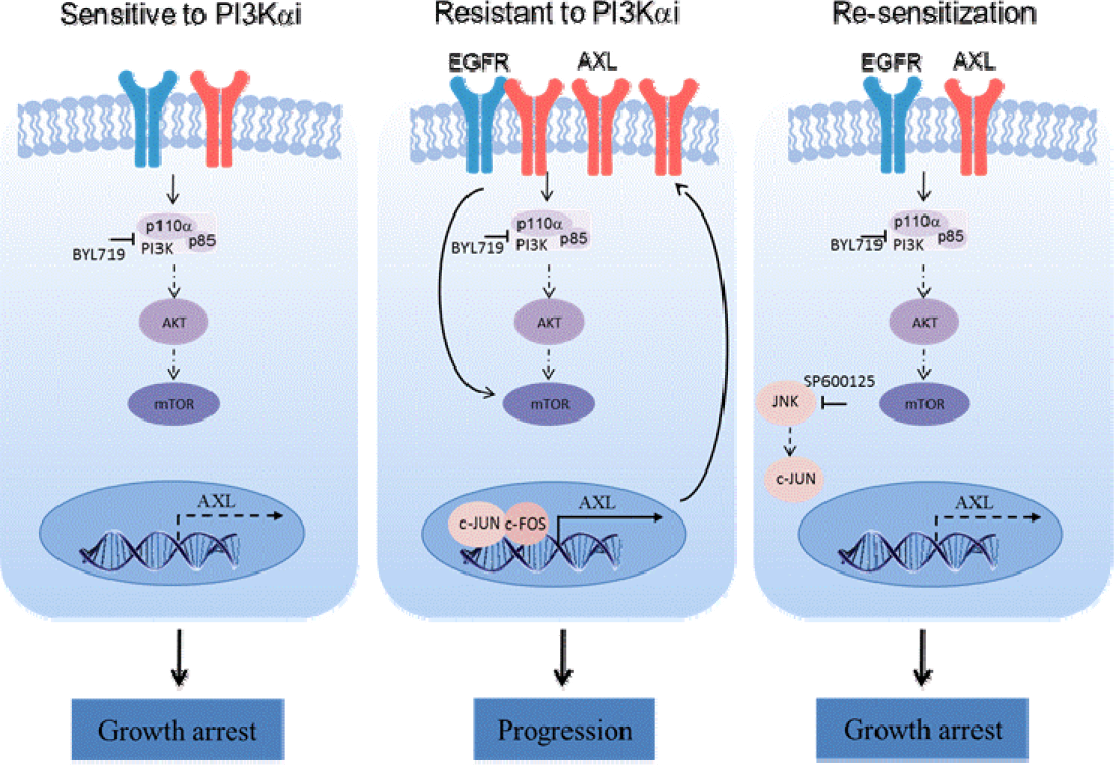
Proposed mechanism of AP-1/ AXL driven resistance to PI3Kαi in HNSCC and ESCC. Cells that acquire resistance to PI3Kαi upregulate AXL via the AP-1 transcription factors c-Jun and c-FOS. AXL dimerizes with EGFR and activates mTOR signaling in an AKT independent manner. Combination of the PI3Kα inhibitor BYL719 with the JNK inhibitor SP600125 sensitizes the cells to BYL719.

Our rationale for targeting the PI3K pathway with BYL719 (an isoform-specific inhibitor) is based on the encouraging results obtained from pre-clinical and clinical studies that tested the efficacy of the p110α inhibitor BYL719 in PIK3CA^mut^ cancers (13, 31–37). The major obstacle to implementing BYL719 therapy in HNSCC and ESCC lies in the primary and acquired resistance to this compound, which is driven by the overexpression of AXL (14). This overexpression of AXL in response to therapy is not unique to these malignant diseases or to PI3K-targeting therapies, as AXL pathway activation has been reported in various cancers that developed resistance to radiotherapy, chemotherapies and EGFR inhibitors (15–19, 38–40). Indeed, small molecule inhibitors (such as BGB324, ONO-747, TP-0903, MGCD516 and BPI-9016M) or specific antibodies (CAB-AXL-ADC) that target the AXL receptor or the AXL pathway are currently under clinical evaluation in 14 clinical trials as a monotherapy or in combination with other therapies in different settings (ClinicalTrials.gov). The clinical outcomes of these studies are not yet available, but pre-clinical models indicate that the efficacy of a single agent targeting AXL signaling can be limited by the upregulation of other receptors, such as MERTK (41), while the potency of several combination therapies is superior to that of the monotherapies (15, 40, 42–47).

Here, we adopted an alternative approach to reducing the availability of AXL by limiting its transcription. Our initial demonstration that AXL levels determine the efficacy of treatment with BYL719 in HNSCC and ESCC cells in vitro and in vivo supported the rationale to reduce AXL expression in tumor cells as a means to enhancing BYL719 efficacy. Our results suggest that this approach could be beneficial when applied to HPV^Neg^ as well as to HPV^Pos^ cells that may be intrinsically resistant to BYL719 (48). Additional confidence in our approach to reduce the AXL expression level as a therapeutic opportunity was provided by the study of Wheeler’s group showing that a reduction of AXL expression enhanced the efficacy of cetuximab in HPV^Pos^ and HPV^Neg^ HNSCC tumors (15, 49).

A number of studies have shown that overexpression of AXL in response to therapeutic stress can evolve through a variety of mechanisms, including transcriptional regulation (15, 23, 24, 50–52), post-translational regulation (53, 54), and expression of miRNA (55) [reviewed in (56)]. In our isogenic tumor cell line of BYL719-sensitive and BYL719-acquired-resistant cells, we observed a marked increase in the AXL transcript and a concomitant increase in members of the AP-1 complex in resistant cells. Thus, we posited that the AP-1 transcription factor complex may be responsible for overexpression of AXL in BYL719-resistant cells. Indeed, knockdown of AP-1 complex members c-FOS and c-JUN in tumor cells resulted in significant reductions in AXL protein and mRNA expression levels in BYL719-sensitive and BYL719-resistant cells.

The mechanism by which BYL719 induces overexpression of c-JUN is not yet clear and requires further investigation. A possible explanation may involve MITF regulation of c-JUN expression. It has been shown that MITF negatively regulates c-JUN expression and is itself suppressed by pro-inflammatory cytokines (57). Thus, in light of recent studies demonstrating that tumor cells secrete pro-inflammatory cytokines, such as TNF α, following anti-PI3K treatment (58), it is tempting to suggest that BYL719 treatment induces the secretion of pro-inflammatory cytokines, which may downregulate MITF expression and hence enhance c-JUN expression.

In an attempt to target c-JUN induced-AXL expression, we used SP600125, an inhibitor of JNK activation, as direct c-JUN inhibitors are not available (29). Supplementation of BYL719 with SP600125 improved the anti-proliferative effect of BYL719 in vitro to a degree similar to that of silencing of c-JUN by using RNAi. Since the combination of BYL719 with SP600125 was shown to have a synergistic anti-proliferative effect in most of the 15 cell lines tested, we used relatively low dosages of BYL719 and SP600125 (25 mg/kg and 15 mg/kg daily, respectively) in the testing of the combination in vivo. These low drug concentrations prevented tumor progression with no evidence of toxicity.

Targeting AP-1 or c-JUN signaling may have a broader effect on tumor cells than simply reducing AXL expression. Targeting the pleotropic role of AP-1 TFs in HNSCC may affect the cancer stem cells (59), epithelial-mesenchymal transition (60) and/or tumor invasiveness (61). Additionally, blocking AP-1 transcription activity may serve to influence the expression of key RTKs, such as EGFR (62, 63), and/or to reduce the expression of the immuno-suppressor modulator of programmed cell death ligand 1 (PD-L1) (64), and hence enhance tumor elimination. The latter possibility is also supported by the findings that AXL and PD-L1 expression are correlated in different cancers (65, 66), and in the cancer genome atlas (TCGA) data set for head and neck cancer (data not shown).

In summary, we present evidence that upregulation of AXL in PI3K-resistant cells is regulated by AP-1 TFs. Knockdown of AXL or AP-1 enhanced the anti-tumor efficacy of BYL719. Targeting of JNK with BYL719 in combination with SP600125 provided potent anti-tumor activity against HPV^Pos^ and HPV^Neg^ cells. These results provide strong grounds for investigating drug combinations of PI3K- and JNK-targeting therapies in HNSCC and ESCC.

## Materials and Methods

### Tumor cell lines

All HPV^Neg^ cell lines were purchased from commercial vendors. Cell line sources and specific media are described in Supplementary Materials and Methods. BYL719-resistant KYSE180^Res^ and CAL33^Res^ cell lines were developed previously (14), and the SNU1076-resistant cell line was developed using the same approach as that described previously (14). All HPV^Pos^ cell lines, which were available in the laboratory of Prof. Reidar Grénman, were grown in DMEM medium. Two mouse HNSCC cell lines were developed in our laboratory from 4NQO-induced oral cancer. Briefly, fresh lip and tongue tumor tissues were washed with Hank’s balanced salt solution (HBSS) or phosphate buffered saline (PBS), cut into small pieces with sterile scissors, and treated with an enzyme mixture of collagenase (10 mg/ml), hyaluronidase (1 mg/ml), and DNase (200 U/ml). The tissues were dissociated using the gentleMACS*™* Dissociator. Cells were filtered through a 70-μM cell strainer, centrifuged (1500 rpm, 5 minutes) and cultured in DMEM medium. The cultures were stained for the epithelial markers cytorkeratin 14 and E-cadherin for verification of epithelial origin.

All cells were maintained at 37°C in a humidified atmosphere at 5% CO_2_, in the relevant media supplemented with 1% L-glutamine 200 mM, 100 units each of penicillin and streptomycin, and 10% fetal bovine serum. Acquired resistance cell lines were grown in the presence of BYL719 (2 μM), which was added to the medium every 3 days. Cells were routinely tested for mycoplasma infection, and treated with appropriate antibiotics as needed (De-Plasma™, TOKU-E, D022). HPV^Neg^ cells were tested for cell line authentication in our previous work (14), and several cell lines (SNU1076, FADU, HSC-2 and HSC-4) were re-tested in this work.

### Western blotting

Cells were washed with ice-cold PBS, harvested into lysis buffer (50 mM HEPES, pH 7.5, 150 mM NaCl, 1 mM EDTA, 1 mM EGTA, 10% glycerol, 1% Triton X-100, 10 μM MgCl_2_), supplemented with phosphatase inhibitor cocktails (Biotool Cat. B15001A/B) and protease inhibitor (Sigma-Aldrich Cat. P2714-1BTL), and placed on ice for 30 minutes, followed by 3 minutes of ultrasonic cell disruption. Lysates were cleared by centrifugation (30 minutes,14,000 rpm, 4°C). Supernatants were collected, protein concentrations were determined using Bio-Rad protein assay, SDS sample buffer was added, and samples were boiled for 5 minutes before being frozen at −80°C until further use. Whole cell lysates (25 μg) were separated on 10% SDS–PAGE and blotted onto PVDF membranes (BioRad trans blot^®^ Turbo™ transfer pack #1704157). Membranes were blocked for 1 hour in blocking solution [5% BSA (Amresco 0332-TAM) in Tris-buffered saline (TBS) with 0.1% Tween] and then incubated with primary antibodies diluted in blocking solution. Mouse and rabbit horseradish peroxidase (HRP)-conjugated secondary antibodies were diluted in blocking solution. Protein-antibody complexes were detected by chemiluminescence [Westar Supernova (Cyanagen Cat. XLS3.0100) and Westar Nova 2.0 (Cyanagen Cat. XLS071.0250)], and images were captured using the Azure C300 Chemiluminescent Western Blot Imaging System, Azure Biosystems. details of antibodies and dilutions used are presented in Supplementary Materials and Methods

### Real-time PCR

Total RNA was isolated from the cultured cells using PureLink™ RNA Mini Kit (Invitrogen, Thermo Fisher Scientific) according to the manufacturer’s protocol, and 1 μg RNA was converted to cDNA using qScript™ cDNA synthesis kit (Quanta Bioscience, 95047-100) according to the manufacturer’s protocol. Real-time PCR was performed (Roche light cycler 480 II) using PrimeTime Gene Expression Master Mix (IDT Cat. 1055770), with matching probes from IDT: Axl gene: Hs.PT.56a.1942285, GAPH gene: Hs.PT.39a.22214836, Jun gene: Hs.PT.58.25094714.g and Fos gene: Hs.PT.58.15540029. Analysis was performed with LightCycler^®^ 480 Software, Version 1.5.1. Fold change was calculated using the ΔΔCt method. Results were normalized to GAPDH levels and presented as a fold increase of the control cells.

### Gene set enrichment analysis

Seven hundred and fifteen genes were upregulated in resistant cells compared to sensitive cells (cut off of >100 reads, and > of 0.5log_2_, padj >0.05). Gene set enrichment analysis was performed using GESA of the Broad Institute (http://software.broadinstitute.org/gsea/index.jsp). The threshold for a conserved motif was defined as (padj>0.1×10^−5^). For the binding site of the AXL promoter, we used the Genecard and the TF binding sites on QIAGEN websites. All the data is available in Supplementary Table 1.

### Immunohistochemistry and immunofluorescence

Tissues were fixed in a 4% paraformaldehyde (PFA) solution for a maximum of 24 hours at room temperature, dehydrated, and embedded in paraffin. The tissue sections were de-paraffinized with xylene. H_2_O_2_, 3%, was used to block the endogenous peroxidase activity for 20 minutes, and thereafter the sections were rinsed in water for 5 minutes. Antigen retrieval was performed in citrate buffer (pH 6) at 99.99°C for 5 minutes. Sections were then blocked for 1 hour at room temperature with blocking solution [PBS, 0.1% Tween (0.0125% for AXL staining), 5% BSA], followed by incubation with primary antibody (diluted in blocking solution) overnight at 4°C. The ABC kit (VECTASTAIN Cat. VE-PK-6200) was used for detection according to the manufacturer’s protocol. Sections were counter-stained with hematoxylin and mounted in mounting medium (Micromount, Leica Cat. 380-1730).

For IF, cells were seeded on 8-well glass slides (Cellvis, Cat no. C8-1.5H-N) for 48 hours. Cells were rinsed with cold PBS (4°C) and fixed in 4% PFA for 30 minutes at room temperature. Thereafter, cells were rinsed with PBS, followed by permeabilization on ice for 5 minutes in PBS with 0.05% Triton X-100 (Sigma-Aldrich), re-rinsed with PBS, and blocked in blocking solution for 30 minutes at room temperature (5% BSA in Tween-PBS). The cells were incubated with primary antibodies overnight at 4°C, rinsed with PBS, and incubated with Cy3 conjugated anti-rabbit secondary antibody (1:250) (Jackson ImmunoResearch, 111-165-144) and Alexa Fluor^®^ 488-conjugated anti-mouse secondary antibody (1:250) (Jackson ImmunoResearch, 115-545-062) at room temperature for 1 hour. Cells were then rinsed with PBS and mounted in DAPI Fluoromount-G^®^ (SouthenBiotech, 0100-20).

IHC and IF slides were digitalized using the Pannoramic Scanner (3DHISTECH, Budapest, Hungary) and analyzed using QuantCenter (3DHISTECH, Budapest, Hungary) using a single threshold parameter for all images of a specific staining in each experiment.

### Tissue microarrays and HNSCC samples

Tissue microarray slides containing 17 HNSCC and 15 benign tumors (in duplicate) were purchased from US Biomax (HNT961). The information regarding these samples and the levels of AXL and c-JUN are summarized in Supplementary Table 2. Fresh HNSCC specimens were collected from Soroka Medical Center Surgey room, and stored in liquid nitrogen. Helsinki was approved by the Soroka Medical Center (#0103-17-SOR and 0421-16-SOR).

### IC:50 and synergy assay

Cells were seeded in 96-well plates (3000 cells per well), treated with increasing concentrations of the relevant drugs (0-10 μM), and allowed to proliferate for 4 days. Cells were then fixed with 0.6 M trichloroacetic acid (TCA) for 1 hour at 4°C, rinsed, and stained with crystal violet (1g/L) for 10 minutes at room temperature. Following additional rinsing, the bound crystal violet was dissolved out with 10% acetic acid, and absorbance was measured at 570 nm (BioTek™ Epoch™ spectrophotometer). IC:50 values were calculated using the GraphPad Software. For the synergy assays, the proliferation of the cells in the different treatment groups was presented as a percentage of control (DMSO treated) cells and the percent of growth inhibition was calculated. Synergy scores were analyzed using Chalice™ Bioinformatics Software (http://cwr.horizondiscovery.com) (67), and the synergistic effects were calculated based on the statistic Loewe Excess model, according to the equation:

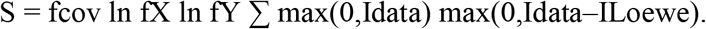

### siRNA and shRNA

For transient silencing of c-JUN and c-FOS, cells were transfected using GenMute^®^ siRNA Transfection Reagent (SignaGen Cat. SL100568) according to the manufacturer’s protocol, with an siRNA non-targeting control sequence (IDT, Cat. 51-01-14-04) and a c-JUN or c-FOS gene targeting sequence (IDT, Cat. hs.Ri.JUN.13 and hs.Ri.FOS.13.1, respectively). Cells were harvested after 48 hours for WB and qPCR analysis. For BYL719 IC:50 experiments, cells were treated with the relevant drugs following 24 hours of transection and allowed to proliferate for an additional 4 days.

For the production of shRNAs cell lines, we created lentiviruses by transfecting HEK293 cells with the viral plasmids psPAX2, pMD2.G, and PLKO with shRNAs – a control scrambled sequence (shCT) or 2 different sequences for the silencing of AXL expression (shAXL1 and shAXL2, MSKCC RNAi core facility identifiers TRCN0000001037 and TRCN0000194771, respectively) using PolyJet transfection reagent (SignaGen, Cat. SL100688) according to the manufacturer’s protocol. Viruses were collected after 48-72 hours and used for cell infection. Cells were seeded in a 6-well plates (150000 cells per well) and infected with the lentiviruses in the presence of Polybrene (Sigma Aldrich Cat. 5G-H9268). Cells were selected with puromycin (Gibco Cat. A11138-03).

### Cell proliferation assay

Cells were seeded in 24-well plates (10,000 cells per well) and treated as indicated in the Results section. The real time cell history recorder JULI™ Stage was used to record cell confluence every 6 hours. Results are presented as the averaged confluence ±SEM. For the 4NQO-induced cell lines (Figure 6B), cells were treated as indicated in Figure 6B for 4 days and then fixed and stained with crystal violet. Results are presented as a percentage of the control (DMSO-treated) cells.

### Establishment of tumor xenografts and studies in mice

NOD.CB17-Prkdc-scid/NCr Hsd (Nod.Scid) and C57/BL/6 mice were purchased from Envigo. NOD.Cg-Prkdc Il2rg/SzJ (NSG) mice were purchased from Jackson labs.

For cell-line-derived xenografts (CDXs), 6-week-old Nod.Scid mice were injected subcutaneously in the flank with 2×10^6^ cells in 200 μl PBS (100 μl in each side). Tumors (60 mm^3^) developed after about 2 weeks.

Patient-derived xenografts (PDXs) were established from HNSCC patients treated in the Ear, Nose, and Throat Unit, Soroka Medical Center, Beer-Sheva, Israel. All patients signed informed consent forms. All PDXs were first transplanted subcutaneously into the flanks of 6-week-old NSG mice. Upon successful tumor engraftment tumor were expanded and re-transplanted into Nod.Scid mice for drug efficacy experiments. About 2 weeks after the second transplantation, the mice were randomized into four groups of 6–8 mice per group.

To produce murine HNSCC, C57BL/6 mice were given 50 μg/mL 4NQO in their drinking water for 3-4 months, and tumors developed 2 months later (Supplementary Figure 6A). The lip and tougher tumor model were expended in secondary mice.

For survival experiments, an orthotropic injection of 0.5×10^6^ 4NQO-tongue cell line cells into the tongues of syngeneic C57BL/6 mice. At a tumor volume of 70 to 120 mm^3^ (2-4 weeks after implantation), animals were randomized into four groups of 10-12 mice per group.

For the in vivo experiments, animals were treated orally with vehicle [corn oil (Sigma Aldrich cat. C8267-500ML) containing 4% DMSO (Sigma Aldrich cat. D8418) for SP600125 administration or 0.5% carboxymethylcellulose (Sigma Aldrich cat. 9481-1KG) for BYL719 administration], BYL719 (25 mg/kg) and/or JNK inhibitor SP600125 (15 mg/kg) daily. Tumors were measured with a digital caliper twice a week, and tumor volumes were determined according to the formula: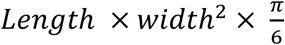. At the end of the experiment, animals were sacrificed by subjecting them to CO_2_ inhalation, and the tumors were harvested for investigation. Tumor volumes were normalized to initial volumes and presented as an averaged percentage of the initial volumes ± SEM.

Mice were maintained and treated according to the institutional guidelines of Ben-Gurion University of the Negev. Mice were housed in air-filtered laminar flow cabinets with a 12 hours light/dark cycle and food and water ad libitum. Animal experiments were approved by the Institutional Animal Care and Use Committee (IL.80-12-2015).

### Drugs

BYL719 was kindly provided by Novartis. For in vitro experiments BYL719 was dissolved in DMSO and for in vivo administration in 0.5% carboxymethylcellulose. SP600125 was purchased from Fluorochem (Cat. M02529). For in vitro assays, SP600125 was dissolved in DMSO and for in vivo administration in corn oil with 4% DMSO.

### Statistical analysis

Experiments were repeated at least 2-3 times, and representative data/images are shown. Statistical analysis was performed using GraphPad Prism software, and results are presented as means ± SEM. For comparisons between groups, P values were calculated. For the different experiments, P values of 0.05, 0.01, or 0.001 were considered statistically significant, as indicated by (*), (**), or (***) on the Figures.

## Author contributions

M.B, M.P, L.C and M.E designed research studies. M.B, M.P, N.B and L.C conducted the experiments and analyzed the data. J.B, A.B.D. and R.G. provided clinical samples and cell lines. M.E., L.C. and M.B. wrote the manuscript.

## Acknowledgments

This work was funded by an Israel Science Foundation (ISF, 700/16) grant to M.E., an INCPM (1901/17) grant to ME, and a Concern Foundation (#7895) grant to M.E. M.E is supported by an Alon Fellowship for outstanding young researchers. M.B is supported by the national-VATA-fellowship for excellent Ph.D students.

## References

1. Wiegand S, Zimmermann A, Wilhelm T, and Werner JA. Survival After Distant Metastasis in Head and Neck Cancer. Anticancer Res. 2015;35(10):5499–502.

2. Ferlay J, Soerjomataram I, Dikshit R, Eser S, Mathers C, Rebelo M, Parkin DM, Forman D, and Bray F. Cancer incidence and mortality worldwide: sources, methods and major patterns in GLOBOCAN 2012. Int J Cancer. 2015;136(5):E359–86.

3. Curado MP, and Hashibe M. Recent changes in the epidemiology of head and neck cancer. Curr Opin Oncol. 2009;21(3):194–200.

4. Marur S, D’Souza G, Westra WH, and Forastiere AA. HPV-associated head and neck cancer: a virus-related cancer epidemic. The Lancet Oncology. 2010;11(8):781–9.

5. Argiris A, Karamouzis MV, Raben D, and Ferris RL. Head and neck cancer. Lancet.2008;371(9625):1695–709.

6. Cancer Genome Atlas N. Comprehensive genomic characterization of head and neck squamous cell carcinomas. Nature. 2015;517(7536):576–82.

7. Seiwert TY, Zuo Z, Keck MK, Khattri A, Pedamallu CS, Stricker T, Brown C, Pugh TJ, Stojanov P, Cho J, et al. Integrative and comparative genomic analysis of HPV-positive and HPV-negative head and neck squamous cell carcinomas. Clinical cancer research: an official journal of the American Association for Cancer Research. 2015;21(3):632–41.

8. Stransky N, Egloff AM, Tward AD, Kostic AD, Cibulskis K, Sivachenko A, Kryukov GV, Lawrence MS, Sougnez C, McKenna A, et al. The mutational landscape of head and neck squamous cell carcinoma. Science. 2011;333(6046):1157–60.

9. Cantley LC. The phosphoinositide 3-kinase pathway. Science. 2002;296(5573):1655–7.

10. Courtney KD, Corcoran RB, and Engelman JA. The PI3K pathway as drug target in human cancer. Journal of clinical oncology: official journal of the American Society of Clinical Oncology. 2010;28(6):1075–83.

11. Engelman JA. Targeting PI3K signalling in cancer: opportunities, challenges and limitations. Nature reviews Cancer. 2009;9(8):550–62.

12. Rodon J, Dienstmann R, Serra V, and Tabernero J. Development of PI3K inhibitors: lessons learned from early clinical trials. Nat Rev Clin Oncol. 2013;10(3):143–53.

13. Juric D, Rodon J, Tabernero J, Janku F, Burris HA, Schellens JHM, Middleton MR, Berlin J, Schuler M, Gil-Martin M, et al. Phosphatidylinositol 3-Kinase alpha-Selective Inhibition With Alpelisib (BYL719) in PIK3CA-Altered Solid Tumors: Results From the First-in-Human Study. Journal of clinical oncology: official journal of the American Society of Clinical Oncology. 2018;36(13):1291–9.

14. Elkabets M, Pazarentzos E, Juric D, Sheng Q, Pelossof RA, Brook S, Benzaken AO, Rodon J, Morse N, Yan JJ, et al. AXL mediates resistance to PI3Kalpha inhibition by activating the EGFR/PKC/mTOR axis in head and neck and esophageal squamous cell carcinomas. Cancer cell. 2015;27(4):533–46.

15. Brand TM, Iida M, Stein AP, Corrigan KL, Braverman CM, Luthar N, Toulany M, Gill PS, Salgia R, Kimple RJ, et al. AXL mediates resistance to cetuximab therapy. Cancer research. 2014;74(18):5152–64.

16. Dufies M, Jacquel A, Belhacene N, Robert G, Cluzeau T, Luciano F, Cassuto JP, Raynaud S, and Auberger P. Mechanisms of AXL overexpression and function in Imatinib-resistant chronic myeloid leukemia cells. Oncotarget. 2011;2(11):874–85.

17. Giles KM, Kalinowski FC, Candy PA, Epis MR, Zhang PM, Redfern AD, Stuart LM, Goodall GJ, and Leedman PJ. Axl mediates acquired resistance of head and neck cancer cells to the epidermal growth factor receptor inhibitor erlotinib. Molecular cancer therapeutics. 2013;12(11):2541–58.

18. Hong J, and Belkhiri A. AXL mediates TRAIL resistance in esophageal adenocarcinoma. Neoplasia.2013;15(3):296–304.

19. Zhang Z, Lee JC, Lin L, Olivas V, Au V, LaFramboise T, Abdel-Rahman M, Wang X, Levine AD, Rho JK, et al. Activation of the AXL kinase causes resistance to EGFR-targeted therapy in lung cancer. Nature genetics. 2012;44(8):852–60.

20. Mudduluru G, and Allgayer H. The human receptor tyrosine kinase Axl gene--promoter characterization and regulation of constitutive expression by Sp1, Sp3 and CpG methylation. Bioscience reports. 2008;28(3):161–76.

21. Mudduluru G, Vajkoczy P, and Allgayer H. Myeloid zinc finger 1 induces migration, invasion, and in vivo metastasis through Axl gene expression in solid cancer. Mol Cancer Res. 2010;8(2):159–69.

22. Mudduluru G, Leupold JH, Stroebel P, and Allgayer H. PMA up-regulates the transcription of Axl by AP-1 transcription factor binding to TRE sequences via the MAPK cascade in leukaemia cells. Biology of the cell / under the auspices of the European Cell Biology Organization. 2010;103(1):21–33.

23. Maurus K, Hufnagel A, Geiger F, Graf S, Berking C, Heinemann A, Paschen A, Kneitz S, Stigloher C, Geissinger E, et al. The AP-1 transcription factor FOSL1 causes melanocyte reprogramming and transformation. Oncogene. 2017;36(36):5110–21.

24. Sayan AE, Stanford R, Vickery R, Grigorenko E, Diesch J, Kulbicki K, Edwards R, Pal R, Greaves P, Jariel-Encontre I, et al. Fra-1 controls motility of bladder cancer cells via transcriptional upregulation of the receptor tyrosine kinase AXL. Oncogene. 2012;31(12):1493–503.

25. Zhen Y, Lee IJ, Finkelman FD, and Shao WH. Targeted inhibition of Axl receptor tyrosine kinase ameliorates anti-GBM-induced lupus-like nephritis. J Autoimmun. 2018.

26. Konieczkowski DJ, Johannessen CM, Abudayyeh O, Kim JW, Cooper ZA, Piris A, Frederick DT, Barzily-Rokni M, Straussman R, Haq R, et al. A melanoma cell state distinction influences sensitivity to MAPK pathway inhibitors. Cancer discovery. 2014;4(7):816–27.

27. Muller J, Krijgsman O, Tsoi J, Robert L, Hugo W, Song C, Kong X, Possik PA, Cornelissen-Steijger PD, Geukes Foppen MH, et al. Low MITF/AXL ratio predicts early resistance to multiple targeted drugs in melanoma. Nat Commun. 2014;55712.

28. Xu H, Jin X, Yuan Y, Deng P, Jiang L, Zeng X, Li XS, Wang ZY, and Chen QM. Prognostic value from integrative analysis of transcription factors c-Jun and Fra-1 in oral squamous cell carcinoma: a multicenter cohort study. Sci Rep. 2017;7(1):7522.

29. Bennett BL, Sasaki DT, Murray BW, O’Leary EC, Sakata ST, Xu W, Leisten JC, Motiwala A, Pierce S, Satoh Y, et al. SP600125, an anthrapyrazolone inhibitor of Jun N-terminal kinase. Proceedings of the National Academy of Sciences of the United States of America. 2001;98(24):13681–6.

30. Hawkins BL, Heniford BW, Ackermann DM, Leonberger M, Martinez SA, and Hendler FJ. 4NQO carcinogenesis: a mouse model of oral cavity squamous cell carcinoma. Head & neck. 1994;16(5):424–32.

31. Rodon J, Juric D, Gonzalez-Angulo A-M, Bendell J, Berlin J, Bootle D, Gravelin K, Huang A, Derti A, Wuerthner JLJ, et al. Annual meeting AACR 2013. 2013.

32. Elkabets M, Vora S, Juric D, Morse N, Mino-Kenudson M, Muranen T, Tao J, Campos AB, Rodon J, Ibrahim YH, et al. mTORC1 Inhibition Is Required for Sensitivity to PI3K p110alpha Inhibitors in PIK3CA-Mutant Breast Cancer. Science translational medicine. 2013;5(196):196ra99.

33. Fritsch C, Huang A, Chatenay-Rivauday C, Schnell C, Reddy A, Liu M, Kauffmann A, Guthy D, Erdmann D, De Pover A, et al. Characterization of the novel and specific PI3Kalpha inhibitor NVP-BYL719 and development of the patient stratification strategy for clinical trials. Molecular cancer therapeutics. 2014;13(5):1117–29.

34. Furet P, Guagnano V, Fairhurst RA, Imbach-Weese P, Bruce I, Knapp M, Fritsch C, Blasco F, Blanz J, Aichholz R, et al. Discovery of NVP-BYL719 a potent and selective phosphatidylinositol-3 kinase alpha inhibitor selected for clinical evaluation. Bioorganic & medicinal chemistry letters. 2013;23(13):3741–8.

35. Zumsteg ZS, Morse N, Krigsfeld G, Gupta G, Higginson DS, Lee NY, Morris L, Ganly I, Shiao SL, Powell SN, et al. Taselisib (GDC-0032), a Potent beta-Sparing Small Molecule Inhibitor of PI3K, Radiosensitizes Head and Neck Squamous Carcinomas Containing Activating PIK3CA Alterations. Clinical cancer research: an official journal of the American Association for Cancer Research. 2016;22(8):2009–19.

36. De Felice F, and Guerrero Urbano T. New drug development in head and neck squamous cell carcinoma: The PI3-K inhibitors. Oral oncology. 2017;67119–23.

37. Mizrachi A, Shamay Y, Shah J, Brook S, Soong J, Rajasekhar VK, Humm JL, Healey JH, Powell SN, Baselga J, et al. Tumour-specific PI3K inhibition via nanoparticle-targeted delivery in head and neck squamous cell carcinoma. Nat Commun. 2017;8(14292).

38. Liu L, Greger J, Shi H, Liu Y, Greshock J, Annan R, Halsey W, Sathe GM, Martin AM, and Gilmer TM. Novel mechanism of lapatinib resistance in HER2-positive breast tumor cells: activation of AXL. Cancer research. 2009;69(17):6871–8.

39. Byers LA, Diao L, Wang J, Saintigny P, Girard L, Peyton M, Shen L, Fan Y, Giri U, Tumula PK, et al. An epithelial-mesenchymal transition gene signature predicts resistance to EGFR and PI3K inhibitors and identifies Axl as a therapeutic target for overcoming EGFR inhibitor resistance. Clinical cancer research: an official journal of the American Association for Cancer Research. 2013;19(1):279–90.

40. Brand TM, Iida M, Stein AP, Corrigan KL, Braverman CM, Coan JP, Pearson HE, Bahrar H, Fowler TL, Bednarz BP, et al. AXL Is a Logical Molecular Target in Head and Neck Squamous Cell Carcinoma. Clinical cancer research: an official journal of the American Association for Cancer Research. 2015;21(11):2601–12.

41. McDaniel NK, Cummings CT, Iida M, Hulse J, Pearson HE, Vasileiadi E, Parker RE, Orbuch RA, Ondracek OJ, Welke NB, et al. MERTK mediates intrinsic and adaptive resistance to AXL-targeting agents. Molecular cancer therapeutics. 2018.

42. Balaji K, Vijayaraghavan S, Diao L, Tong P, Fan Y, Carey JP, Bui TN, Warner S, Heymach JV, Hunt KK, et al. AXL Inhibition Suppresses the DNA Damage Response and Sensitizes Cells to PARP Inhibition in Multiple Cancers. Mol Cancer Res. 2017;15(1):45–58.

43. Ben-Batalla I, Erdmann R, Jorgensen H, Mitchell R, Ernst T, von Amsberg G, Schafhausen P, Velthaus JL, Rankin S, Clark RE, et al. Axl Blockade by BGB324 Inhibits BCR-ABL Tyrosine Kinase Inhibitor Sensitive and -Resistant Chronic Myeloid Leukemia. Clinical cancer research: an official journal of the American Association for Cancer Research. 2017;23(9):2289–300.

44. Boshuizen J, Koopman LA, Krijgsman O, Shahrabi A, van den Heuvel EG, Ligtenberg MA, Vredevoogd DW, Kemper K, Kuilman T, Song JY, et al. Cooperative targeting of melanoma heterogeneity with an AXL antibody-drug conjugate and BRAF/MEK inhibitors. Nat Med. 2018;24(2):203–12.

45. Guo G, Gong K, Ali S, Ali N, Shallwani S, Hatanpaa KJ, Pan E, Mickey B, Burma S, Wang DH, et al. A TNF-JNK-Axl-ERK signaling axis mediates primary resistance to EGFR inhibition in glioblastoma. Nat Neurosci. 2017;20(8):1074–84.

46. Palisoul ML, Quinn JM, Schepers E, Hagemann IS, Guo L, Reger K, Hagemann AR, McCourt CK, Thaker PH, Powell MA, et al. Inhibition of the Receptor Tyrosine Kinase AXL Restores Paclitaxel Chemosensitivity in Uterine Serous Cancer. Molecular cancer therapeutics. 2017;16(12):2881–91.

47. Sen T, Tong P, Diao L, Li L, Fan Y, Hoff J, Heymach JV, Wang J, and Byers LA. Targeting AXL and mTOR Pathway Overcomes Primary and Acquired Resistance to WEE1 Inhibition in Small-Cell Lung Cancer. Clinical cancer research: an official journal of the American Association for Cancer Research. 2017;23(20):6239–53.

48. Brand TM, Hartmann S, Bhola NE, Li H, Zeng Y, O’Keefe RA, Ranall MV, Bandyopadhyay S, Soucheray M, Krogan NJ, et al. Cross-talk Signaling between HER3 and HPV16 E6 and E7 Mediates Resistance to PI3K Inhibitors in Head and Neck Cancer. Cancer research. 2018;78(9):2383–95.

49. Leonard B, Brand TM, O’Keefe RA, Lee ED, Zeng Y, Kemmer JD, Li H, Grandis JR, and Bhola NE. BET Inhibition Overcomes Receptor Tyrosine Kinase-Mediated Cetuximab Resistance in HNSCC. Cancer research. 2018;78(15):4331–43.

50. Li M, Lu J, Zhang F, Li H, Zhang B, Wu X, Tan Z, Zhang L, Gao G, Mu J, et al. Yes-associated protein 1 (YAP1) promotes human gallbladder tumor growth via activation of the AXL/MAPK pathway. Cancer letters. 2014;355(2):201–9.

51. Ghiso E, Migliore C, Ciciriello V, Morando E, Petrelli A, Corso S, De Luca E, Gatti G, Volante M, and Giordano S. YAP-Dependent AXL Overexpression Mediates Resistance to EGFR Inhibitors in NSCLC. Neoplasia. 2017;19(12):1012–21.

52. Boregowda RK, Medina DJ, Markert E, Bryan MA, Chen W, Chen S, Rabkin A, Vido MJ, Gunderson SI, Chekmareva M, et al. The transcription factor RUNX2 regulates receptor tyrosine kinase expression in melanoma. Oncotarget. 2016;7(20):29689–707.

53. Bae SY, Hong JY, Lee HJ, Park HJ, and Lee SK. Targeting the degradation of AXL receptor tyrosine kinase to overcome resistance in gefitinib-resistant non-small cell lung cancer. Oncotarget. 2015;6(12):10146–60.

54. Miller MA, Oudin MJ, Sullivan RJ, Wang SJ, Meyer AS, Im H, Frederick DT, Tadros J, Griffith LG, Lee H, et al. Reduced Proteolytic Shedding of Receptor Tyrosine Kinases Is a Post-Translational Mechanism of Kinase Inhibitor Resistance. Cancer discovery. 2016;6(4):382–99.

55. Yun MR, Lim SM, Kim SK, Choi HM, Pyo KH, Kim SK, Lee JM, Lee YW, Choi JW, Kim HR, et al. Enhancer Remodeling and MicroRNA Alterations Are Associated with Acquired Resistance to ALK Inhibitors. Cancer research. 2018;78(12):3350–62.

56. Scaltriti M, Elkabets M, and Baselga J. Molecular Pathways: AXL, a Membrane Receptor Mediator of Resistance to Therapy. Clinical cancer research: an official journal of the American Association for Cancer Research. 2016;22(6):1313–7.

57. Riesenberg S, Groetchen A, Siddaway R, Bald T, Reinhardt J, Smorra D, Kohlmeyer J, Renn M, Phung B, Aymans P, et al. MITF and c-Jun antagonism interconnects melanoma dedifferentiation with pro-inflammatory cytokine responsiveness and myeloid cell recruitment. Nat Commun. 2015;68755.

58. Usman MW, Gao J, Zheng T, Rui C, Li T, Bian X, Cheng H, Liu P, and Luo F. Macrophages confer resistance to PI3K inhibitor GDC-0941 in breast cancer through the activation of NF-kappaB signaling. Cell death & disease. 2018;9(8):809.

59. Shrivastava S, Steele R, Sowadski M, Crawford SE, Varvares M, and Ray RB. Identification of molecular signature of head and neck cancer stem-like cells. Sci Rep. 2015;5(7819.

60. Bakiri L, Macho-Maschler S, Custic I, Niemiec J, Guio-Carrion A, Hasenfuss SC, Eger A, Muller M, Beug H, and Wagner EF. Fra-1/AP-1 induces EMT in mammary epithelial cells by modulating Zeb1/2 and TGFbeta expression. Cell Death Differ. 2015;22(2):336–50.

61. Dong C, Ye DX, Zhang WB, Pan HY, Zhang ZY, and Zhang L. Overexpression of c-fos promotes cell invasion and migration via CD44 pathway in oral squamous cell carcinoma. J Oral Pathol Med. 2015;44(5):353–60.

62. Zenz R, Scheuch H, Martin P, Frank C, Eferl R, Kenner L, Sibilia M, and Wagner EF. c-Jun regulates eyelid closure and skin tumor development through EGFR signaling. Dev Cell. 2003;4(6):879–89.

63. Fang Y, Wang Y, Wang Y, Meng Y, Zhu J, Jin H, Li J, Zhang D, Yu Y, Wu XR, et al. A new tumour suppression mechanism by p27Kip1: EGFR down-regulation mediated by JNK/c-Jun pathway inhibition. Biochem J. 2014;463(3):383–92.

64. Green MR, Rodig S, Juszczynski P, Ouyang J, Sinha P, O’Donnell E, Neuberg D, and Shipp MA. Constitutive AP-1 activity and EBV infection induce PD-L1 in Hodgkin lymphomas and posttransplant lymphoproliferative disorders: implications for targeted therapy. Clinical cancer research: an official journal of the American Association for Cancer Research. 2012;18(6):1611–8.

65. Skinner HD, Giri U, Yang LP, Kumar M, Liu Y, Story MD, Pickering CR, Byers LA, Williams MD, Wang J, et al. Integrative Analysis Identifies a Novel AXL-PI3 Kinase-PD-L1 Signaling Axis Associated with Radiation Resistance in Head and Neck Cancer. Clinical cancer research: an official journal of the American Association for Cancer Research. 2017;23(11):2713–22.

66. Aguilera TA, Rafat M, Castellini L, Shehade H, Kariolis MS, Hui AB, Stehr H, von Eyben R, Jiang D, Ellies LG, et al. Reprogramming the immunological microenvironment through radiation and targeting Axl. Nat Commun. 2016;7(13898.

67. Hart LS, Rader J, Raman P, Batra V, Russell MR, Tsang M, Gagliardi M, Chen L, Martinez D, Li Y, et al. Preclinical Therapeutic Synergy of MEK1/2 and CDK4/6 Inhibition in Neuroblastoma. Clinical cancer research: an official journal of the American Association for Cancer Research. 2017;23(7):1785–96.

